# Assessing toxicity and competitive fitness of vibrios isolated from coastal waters in Israel

**DOI:** 10.1101/2025.01.15.632581

**Authors:** Katarzyna Kanarek, Kinga Keppel, Hadar Cohen, Chaya Mushka Fridman, Motti Gerlic, Dor Salomon

## Abstract

Several species of aquatic bacteria belonging to the genus *Vibrio* are emerging pathogens of humans and marine animals. *Vibrio*-associated infections have been shown to correlate with the increase in the oceans’ surface water temperatures. A slow yet steady increase in vibrios isolated from clinical settings in Israel over the past decade led us to investigate the pathogenic potential of vibrios in Israel’s coastal waters. We isolated and sequenced the genomes of 23 vibrios from the Mediterranean and the Red Sea. Analysis of these genomes revealed the presence of diverse toxin secretion systems and toxins, as well as mobile genetic elements known to facilitate the dissemination of fitness-enhancing determinants. Moreover, we showed that at least 10 of these isolates can induce cell death in bone marrow-derived macrophages and that at least 12 isolates can intoxicate a rival *Vibrio* strain in interbacterial competition. Lastly, we determined the susceptibility profiles of these isolates to common antibiotics used to treat *Vibrio* infections and found widespread resistance to azithromycin. Taken together, our results reveal pathogenic potential within the *Vibrio* population of Israel’s coastal waters and underline the need for continued environmental monitoring of emerging pathogens.

**Importance:** The oceans’ surface water temperatures have increased in past decades due to climate change. This increase correlates with the spread of vibrios, a genus of aquatic bacteria, many of which are pathogens of humans and marine animals. Since *Vibrio*-associated illnesses are rising in Israel, we set out to investigate the *Vibrio* population in Israel’s coastal environments and monitor their pathogenic potential. We found diverse repertoires of predicted toxins, the ability to kill mammalian immune cells, and traits that enhance bacterial fitness, such as antibacterial toxicity and resistance to antibiotics commonly used to treat *Vibrio* infections. These findings indicate that pathogenic traits are circulating within the environmental *Vibrio* population in Israel’s coastal waters and suggest that continued monitoring is essential to identify emerging pathogenic strains.

## Observation

*Vibrio*, a genus of aquatic gram-negative bacteria that inhabit marine and estuarine environments, includes several species pathogenic to humans and marine animals (*e*.*g*., *cholerae, vulnificus, parahaemolyticus*, and *alginolyticus*) (1). *Vibrio*-associated human infections, termed vibriosis, manifest as gastroenteritis and wound or ear infections resulting from the consumption or handling of raw seafood and from recreational swimming (1). Notably, infections are prevalent during warmer months when vibrios thrive in the environment (2). Recent reports highlighted a correlation between climate change-associated elevated temperatures of sea surface water, the spread of pathogenic *Vibrio* species, and an increase in *Vibrio*-associated disease occurrence (2–4).

Annual reports from the Central Laboratories in the Israel Ministry of Health, which are tasked with microbial pathogens surveillance, reveal a slow yet steady increase in the number of vibrios isolated from clinical settings in Israel over the last decade, primarily *V. cholerae* non-O1/O139 and *V. alginolyticus* isolated from ear and blood samples (5). Although research focusing on clinical and environmental *V. vulnificus* in Israel was conducted following a 1996 outbreak (6), the environmental *Vibrio* population in Israel coastal waters, a potential reservoir of pathogenic strains, remains understudied.

In the current study, we set out to investigate the toxic and competitive fitness potential of environmental vibrios isolated in Israel. To this end, we collected coastal water samples from two sites in the Mediterranean (Tel Aviv-Yafo and Ma’agan Michael) and one site in the Red Sea (Eilat) during the summer of 2023, and recovered *Vibrio* colonies growing on selective TCBS media plates (Fig. S1 and Supplementary Methods). Of these, 23 colonies with diverse morphologies were selected for whole-genome sequencing (Table S1). Average Nucleotide Identity (ANI) analyses revealed that the isolates belong to various *Vibrio* species, including *alginolyticus, vulnificus, parahaemolyticus, campbellii, fortis, chagasii, owensii, mediterranii, diabolicus*, and *harveyi*. One isolate, TBV020, had an ANI score below 95% compared to the genomes available in the JSpeciesWS genome database, suggesting that it represents a new species (7); we named it *V. sessaei* in honor of our late colleague, Prof. Guido Sessa.

Various virulence factors have been described in the *Vibrio* pan-genome, amongst them type III secretion system (T3SS) (8), type VI secretion system (T6SS) (9–11), and diverse arsenals of secreted toxins (*e*.*g*., MARTX toxins (12) and Cholera toxin (13)) (1). Notably, many of these virulence factors are encoded on mobile genetic elements (MGEs) shown to be horizontally transferred between vibrios, thus posing a risk for the emergence of new pathogens (14).

We analyzed the genome sequence of the 23 isolates for potential virulence factors, focusing on T3SSs, T6SSs, and MARTX toxins, each of which can simultaneously deliver a repertoire of toxic activities into a host cell (Fig. 1a). Ten isolates, belonging to the species *alginolyticus, campbellii, harveyi, diabolicus*, and *parahaemolyticus*, harbor a T3SS similar to the well-studied T3SS1 from *V. parahaemolyticus* (15) (Fig. S2). The *parahaemolyticus* TBV028 isolate also harbors a T3SS2 (Fig. S3) previously shown to induce gastroenteritis (16, 17). In addition, the two *vulnificus* isolates harbor an MARTX toxin. Of the five T6SS types that have been characterized in vibrios, we identified gene clusters representing four types (Fig. S4 and Dataset S1). Gene clusters comparable to *V. parahaemolyticus* T6SS1, T6SS2, and T6SS4 (18) are found in twelve, fourteen, and one isolates, respectively; clusters comparable to T6SS2 of *V. coralliilyticus* (9) are found in twelve isolates. The latter was recently shown to target eukaryotes (9), whereas the *V. parahaemolyticus* T6SS2 and T6SS4 were predicted to target bacteria (18, 19); *V. parahaemolyticus* T6SS1-like systems were shown to deliver both anti-eukaryotic and antibacterial toxins (20, 21).

**Fig. 1.**
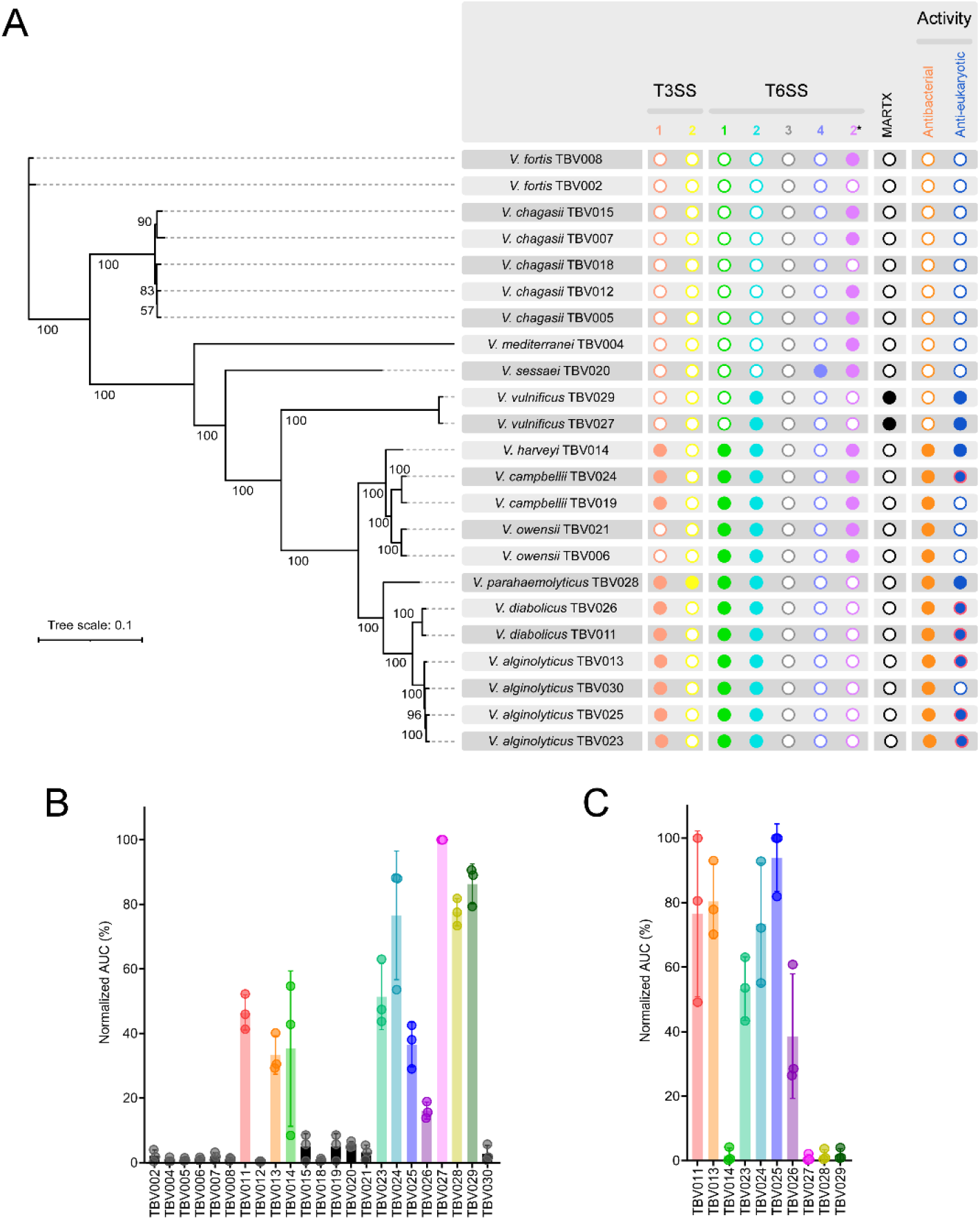
Environmental *Vibrio* isolates harbor toxin delivery systems and induce cell death in macrophages. **(A)** A summary of T3SSs, T6SSs, and MARTX toxins found in the genomes of the 23 indicated isolates. A full circle denotes the presence of the system or toxin. The ability of each isolate to induce cell death in macrophages (anti-eukaryotic; red circles denote contact-independent toxicity) or to intoxicate a competing *V. natriegens* bacterium (antibacterial) is shown on the right. In T6SS, numbers 1-4 denote similarity to T6SS1-4 from *V. parahaemolyticus*, whereas 2* denotes similarity to T6SS2 from *V. coralliilyticus*. The phylogenetic tree is based on the alignment of 2,296 core proteins found in the indicated isolates. The evolutionary history was inferred using the maximum likelihood method. Bootstrap values appear next to the corresponding branch as percent of 100 replicates. **(B-C)** Assessment of cell death upon infection of bone marrow-derived macrophages (BMDMs). Approximately 3.5×10^4^ BMDMs were seeded into 96-well plates in triplicate and infected with the indicated *Vibrio* isolates (B) or with the cleared media of overnight-grown isolates (C). Propidium iodide (PI) was added to the medium prior to infection, and its kinetic uptake was monitored for 5 hours using real-time microscopy (IncucyteZOOM). Cell death was determined as the area under the curve (AUC) of the percentage of PI-positive cells normalized to the number of cells in the wells. Results are shown as the mean ± SD of three independent experiments. An isolate was considered toxic when AUC > 15%.

To investigate the toxic potential of the 23 isolates, we determined their ability to intoxicate murine bone marrow-derived macrophages (BMDMs), serving as model eukaryotic immune cells. We monitored cell death kinetics using real-time microscopy and found that ten of the 23 isolates are detrimental to BMDMs within 5 hours of infection (Fig. 1b and Fig. S9). The toxicity appears to be contact-independent for at least six of these isolates, as evidenced by cell death observed when BMDMs were incubated with cleared media in which bacterial cultures were grown overnight (Fig. 1c and Fig. S10). These results suggest that a secreted toxin is responsible for the toxic phenotype of these six isolates. Surprisingly, the presence of the anti-eukaryotic T3SS1 did not coincide with cell death, as demonstrated by isolates TBV019 and TBV030. Moreover, although the presence of a MARTX toxin in the two *vulnificus* isolates is predicted to induce contact-independent cell death, no contact-independent cell death was observed. Several factors can account for these discrepancies: (i) the growth or infection conditions used in our assays do not induce expression of the relevant systems; (ii) some isolates require longer than 5 hours to induce cell death; (iii) BMDMs are not an appropriate target cell for some isolates.

Next, we sought to assess the competitive fitness of the isolates, since the ability to outcompete rival bacteria was demonstrated to indirectly contribute to virulence (11). When co-incubated on solid agar plates, thirteen isolates either killed or impaired the growth of another marine bacterium, *V. natriegens*, compared to a *V. parahaemolyticus* RIMD 2210633 strain in which the two antibacterial T6SSs were inactivated (Fig. 2a) (19). Notably, this antibacterial toxicity coincided with the presence of a T6SS1-like cluster (Fig. 1a), which was previously shown to be active in vibrios under the assay conditions (i.e., 3% [w/v] NaCl in the media, at 30°C) (9, 19). Furthermore, we recently described MGEs, GMT islands, that are prevalent in vibrios where they serve as mobile armories containing antibacterial T6SS toxins and anti-phage defense systems that enhance competitive fitness and protect against bacteriophage predation, respectively (22). Remarkably, we identified ten GMT islands distributed within six of the 23 isolates (Fig. 2b). Isolate TBV006 harbors four GMT islands, two of which are identical although they are located in different syntenies (Fig. S11). Analysis of the islands’ cargo revealed diverse antibacterial and anti-phage arsenals, confirming our previous findings and indicating that these mobile armories can disseminate within the local *Vibrio* populations to enhance competitive fitness.

**Fig. 2.**
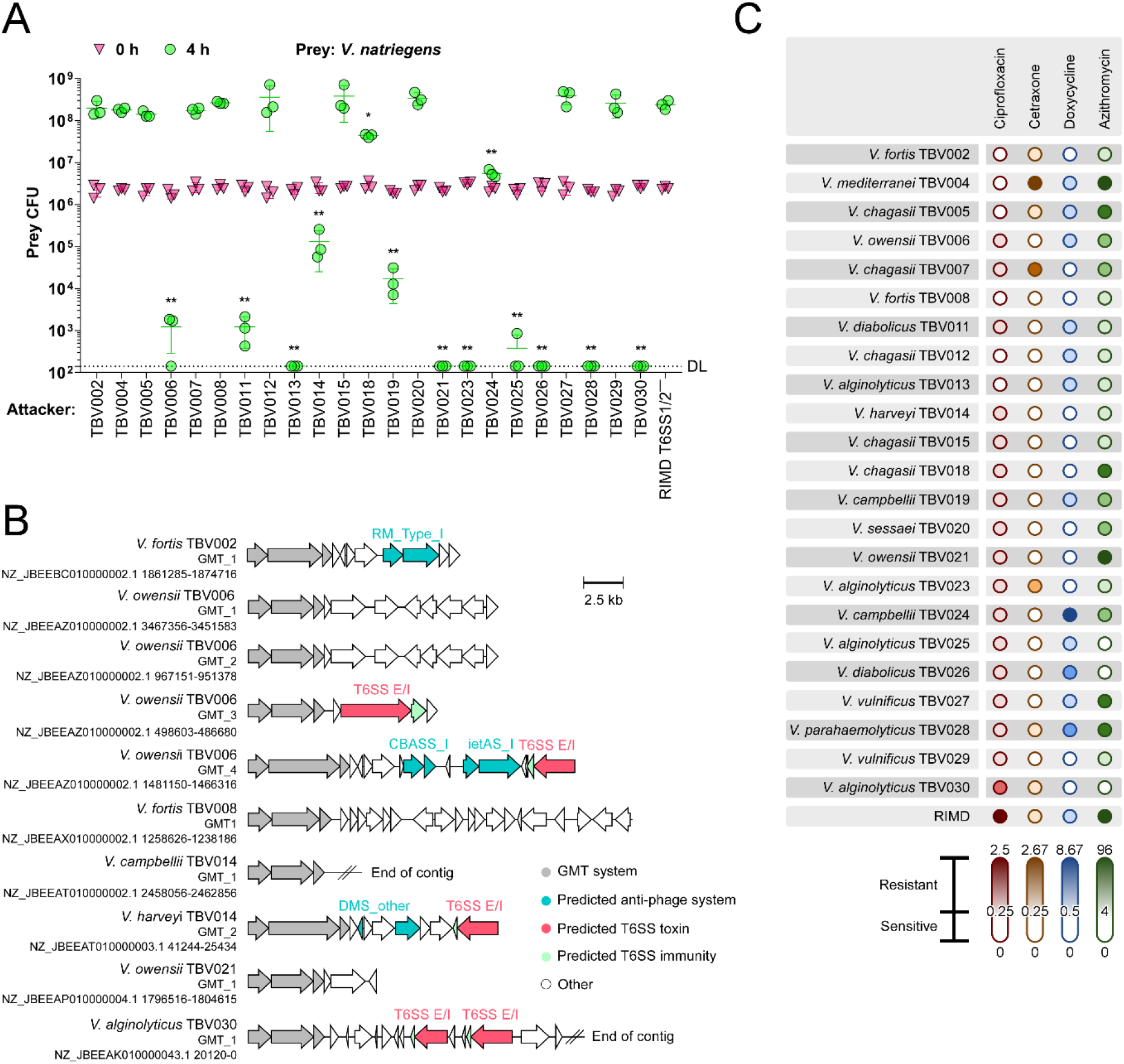
Antibacterial activity and antibiotic resistance in environmental *Vibrio* isolates. **(A)** Viability counts (Colony Forming Units; CFU) of *V. natriegens* prey strains before (0 h) and after (4 h) co-incubation with the indicated *Vibrio* isolate attackers on MLB plates at 30°C. The statistical significance between samples at the 4 h time point was calculated using one-way ANOVA with Dunnett multiple comparisons test on log-transformed data, compared to a *V. parahaemolyticus* RIMD 2210633 strain in which the two T6SSs were inactivated (RIMD T6SS1/2^−^). *, *P* = 0.007; **, *P* < 0.0001. Data are shown as the mean ± SD of 3 biological samples; a representative experiment out of 3 independent experiments is shown. DL, the assay’s detection limit. **(B)** The gene structure and content of GMT islands found in the environmental isolates. RefSeq accessions are denoted. **(C)** The minimum inhibitory concentrations (MICs) of ciprofloxacin, ceftriaxone, doxycycline, and azithromycin were determined for each environmental isolate and for the clinical *V. parahaemolyticus* RIMD 2210633 (RIMD) strain using ETEST® strips. MIC values are displayed as color gradients to represent the resistance levels of each isolate. White color denotes susceptibility; increasingly darker shades denote resistance and higher MIC values. Sensitivity thresholds were set based on EUCAST guidelines, with the following ranges defining susceptibility: 0-0.25 µg/ml for ciprofloxacin and ceftriaxone, 0-0.5 µg/ml for doxycycline, and 0-4 µg/ml for azithromycin. The data shown are the mean of 3 independent experiments.

Lastly, we assessed the susceptibility of the isolates to four antibiotics commonly used to treat vibriosis: ciprofloxacin, ceftriaxone, doxycycline, and azithromycin (Fig. 2c and Dataset S2) (1). The results revealed diverse resistance patterns with no single strain being sensitive or strongly resistant to all antibiotics, although over half of the isolates display strong resistance to at least one drug. Notably, resistance to azithromycin was particularly prevalent, evidenced in sixteen isolates.

## Conclusions

In this study, we isolated 23 *Vibrio* strains from Israel’s coastal waters and demonstrated that they wield toxic activities that can enable them to infect eukaryotic hosts. Moreover, we showed that many of the isolates possess antibacterial determinants that can enhance their competitive fitness in the environment, as well as indirectly contribute to their ability to colonize a host. Furthermore, we observed resistance to several clinically relevant antibiotics within this sample population. Taken together, our results demonstrate pathogenic potential within the *Vibrio* population of Israel’s coastal waters and underline the need to continue surveying these habitats for the emergence of new pathogens.

## Supporting information

Supplementary Methods, Figures, Tables, and References

## Acknowledgements

We thank Udi Qimron for collecting water samples from the Tel Aviv coast. This project received funding from the Israel Science Foundation (ISF grant number 1362/21 to D Salomon and 2174/22 to M Gerlic) and from the Tel Aviv University Center for Combatting Pandemics (to D Salomon). CM Fridman was supported by a scholarship from the Clore Israel Foundation and by a scholarship for outstanding doctoral students from the Orthodox community from the Council for Higher Education. K Kanarek was supported by a PhD Scholarship from the Tel Aviv University Center for Combatting Pandemics. This work was performed in partial fulfillment of the requirements for a PhD degree for Katarzyna Kanarek at the Faculty of Medical and Health Sciences, Tel Aviv University.

## Author contributions

Conceptualization: K Kanarek and D Salomon; Formal Analysis: K Kanarek, H Cohen, CM Fridman, and D Salomon; Funding Acquisition: M Gerlic and D Salomon; Investigation: K Kanarek, K Keppel, CM Fridman, H Cohen, and D Salomon; Methodology: K Kanarek, K Keppel, CM Fridman, and H Cohen; Supervision: D Salomon; Writing – Original Draft Preparation: K Kanarek and D Salomon; Writing – Review and Editing: CM Fridman, K Keppel, H Cohen, and M Gerlic.

