## Supplementary Methods, Figures, Tables, and References for "Assessing toxicity and competitive fitness of vibrios isolated from coastal waters in Israel"

**Supplementary Materials and Methods**

**Supplementary Figure S1-S11**

**Supplementary Table S1**

**Supplementary Dataset S1-S2 (captions)**

**Supplementary References**

### Supplementary Materials and Methods

**Strains and Media:** *Vibrio parahaemolyticus* RIMD 2210633 and its Δ*hcp1*/*hcp2* (T6SS1/2⁻) (1) derivative, *V. natriegens* ATCC 14048, and environmental *Vibrio* isolates (Table S1) were routinely grown in Lysogeny broth containing 3% (w/v) NaCl (MLB, Marine LB) and on Marine Minimal Media (MMM) agar plates (1.5% (w/v) agar, 2% (w/v) NaCl, 0.4% (w/v) galactose, 5 mM $MgSO_4$, 7 mM $K_2SO_4$, 77 mM $K_2HPO_4$, 35 mM $KH_2PO_4$ and 2 mM $NH_4Cl$) at 30°C.

**Isolation of Bacteria:** Seawater samples were collected from three coastal locations in Israel: Tel Baruch Beach in Tel Aviv-Yafo on May 31 and July 5, 2023; Coral Beach in Eilat on June 28, 2023; and Ma'agan Michael on July 16, 2023 (Fig. S1). Water samples were plated onto TCBS agar (Millipore, #86348) and incubated at 30°C for 16 hours. Individual colonies were then transferred to fresh TCBS plates and incubated at 30°C for 16 hours. A total of 23 colonies displaying unique morphologies were selected for further analysis.

**Genomic DNA Extraction and Sequencing:** Genomic DNA (gDNA) was extracted from 23 environmental isolates using the Presto Mini gDNA Bacteria Kit (Geneaid), following the manufacturer's instructions. The gDNA was sequenced with Oxford Nanopore Technology using long-read sequencing technology by Plasmidsaurus (https://www.plasmidsaurus.com/), including construction of an amplification-free long-read sequencing library using v14 library prep chemistry followed by library sequencing using a primer-free protocol. The sequences were then analyzed with the Plasmidsaurus custom analysis and annotation pipeline, including genome assembly and genome annotation using Bakta (2). All genome assemblies were deposited to NCBI under BioProject PRJNA1095381 (Table S1). Isolates were classified into a species based on the highest Average Nucleotide Identity (ANI) scores obtained using the JSpeciesWS online server (https://jspecies.ribohost.com/jspeciesws/); if no known species yielded an ANI score > 95%, the isolate was designated as a new species (i.e., *V. sessaei* TBV020).

**BMDM Infection Assays:** Bone marrow cells from 6- to 8-week-old mice were isolated, and bone marrow-derived macrophages (BMDMs) were obtained after a 7-day differentiation, as previously described (3). Approximately $3.5 \times 10^4$ BMDMs were seeded into 96-well plates in triplicates in 100 µl of FBS and penicillin-streptomycin-free DMEM media.

For infection with live bacteria, *Vibrio* isolates were grown overnight in MLB and added to the wells at an initial $OD_{600}$ = 0.016. The plates were centrifuged for 5 minutes at 400 × *g* to begin the assay.

To monitor contact-independent cell death, *Vibrio* isolates were grown overnight in MLB; in the morning, the cultures were diluted 1:200 in fresh media and incubated for three hours at 30°C. Culture volumes equivalent to 1.0 $OD_{600}$ units were collected in fresh tubes, and DMEM was added to a total volume of 1.0 ml. The bacteria were pelleted by centrifugation, and the supernatants were cleared of bacteria using a 0.2 µm filter. Cleared supernatants were added to the BMDM wells in triplicates (50 µl per well), and the plates were centrifuged for 5 minutes at 400 × *g* to begin the assay.

Propidium iodide (PI; 1.0 µg/ml) was added to the media 30 minutes prior to infection, and its uptake kinetics were assessed every 20 minutes using real-time microscopy (Incucyte SX5) during a 5-hour incubation at 37°C. The data were analyzed using the Incucyte SX5 analysis software and exported to Graphpad PRISM V9. Normalization was performed according to the PI-positive object count to calculate the percentage of dead cells (3). If all cells were stained with PI before the experiment endpoint, the maximum count of PI-positive objects was recorded from that time forward.

**Bacterial Competition Assays:** Bacterial competitions were performed as described previously (4). Briefly, environmental isolate attacker strains were grown overnight in MLB media. *V. natriegens* prey strains containing a plasmid conferring chloramphenicol resistance for selection were grown overnight in MLB media supplemented with 10 μg/ml chloramphenicol. Bacterial cultures were normalized to an $OD_{600}$ = 0.5 and mixed at a 4:1 (attacker:prey) ratio in triplicate. The mixtures were spotted onto MLB agar plates and incubated for 4 hours at 30°C. The viability of the prey strain was determined as colony-forming units growing on selective MLB plates supplemented with 10 μg/ml chloramphenicol at the 0 and 4-hour time points. Statistical analyses and data visualization were performed using GraphPad PRISM v9.

**Assessing Antibiotic Susceptibility:** Minimum Inhibitory Concentrations (MICs) were determined on MLB agar plates overlayed with soft MLB agar (0.5% [w/v]), using the ETEST® method (Biomeriueux), following the manufacturer's instructions. Briefly, *Vibrio* isolates were grown in MLB at 30°C for 16 hours with constant shaking. Each bacterial culture was then diluted 10-fold and 100-fold in fresh media, and the dilution culture that reached an $OD_{600}$ of ~0.9 within 1-2 hours of growth at 30°C was used for subsequent steps. Volumes of refreshed cultures corresponding to $OD_{600}$ units of 0.2 or 1.0 were mixed with 6 ml of molten soft MLB agar and overlaid onto MLB agar plates. The plates were allowed to dry for 1.5 hours before placing ETEST strips on the soft agar overlay (412311 ETEST® Ciprofloxacin (CI), 412303 ETEST® Ceftriaxone (TX), 412328 ETEST® Doxycycline (DC), and 412257 ETEST® Azithromycin (AZ)). Plates were then sealed with parafilm, incubated lid-up at 30°C for 16 hours, and the MICs were recorded according to the border of bacterial growth around the ETEST strip. The assay was performed three times for each bacterial strain. Susceptibility and resistance were determined according to the EUCAST Clinical Breakpoint Tables v. 14.0 for *Vibrio* spp. (January 1, 2024). For ciprofloxacin and ceftriaxone, strains were considered susceptible when MIC ≤ 0.25 mg/l and resistant when MIC > 0.25 mg/l. For doxycycline, strains were considered susceptible when MIC ≤ 0.5 mg/l and resistant when MIC > 0.5 mg/l. For azithromycin, strains were considered susceptible when MIC ≤ 4 mg/l and resistant when MIC > 4 mg/l.

**Constructing a Phylogenetic Species Tree:** The tree was prepared using the M1CR0B1AL1Z3R web server (https://microbializer.tau.ac.il/) (5), with the FASTA genome sequences of the 23 isolates as input. The default parameters were used, except: (i) the minimum protein sequence identity for homologs detection was set to 30%, and (ii) bootstrap analysis was applied to the species tree. In brief, M1CR0B1AL1Z3R computed the core-proteome of the 23 isolates (2,296 proteins) and provided a core-proteome alignment. This alignment was used to construct a maximum-likelihood-based phylogenetic tree using IQ-TREE (6). The resulting tree was visualized using iTOL (https://itol.embl.de) (7).

**Constructing a TssB Phylogenetic Tree:** TssB1-4 from *V. parahaemolyticus* (WP_005480589.1, WP_005463809.1, WP_029843231.1, and WP_042762279.1, respectively) and TssB2 from *V. coralliilyticus* (WP_006962089.1) were used as BLAST queries to identify TssB protein sequences in the genomes of the 23 environmental *Vibrio* isolates (Dataset S2). Unique TssB sequences were aligned using CLUSTAL Omega (8). The evolutionary history was inferred using the Neighbor-Joining method (9). The analysis of TssB included 44 amino acid sequences and 25 conserved sites. The analysis was performed in MEGA11 (10).

**Identifying Secretion Systems and MARTX Toxins:** To identify type III secretion system (T3SS) and type VI secretion system (T6SS) gene clusters, representative T3SS1 and T3SS2β clusters from *V. parahaemolyticus* RIMD 210633 (11) and MAVP-R (12) (respectively), T6SS1 and T6SS2 clusters from *V. parahaemolyticus* RIMD 210633 (13), T6SS3 from *V. parahaemolyticus* 10290 isolate VP160417 (13), T6SS4 from *V. parahaemolyticus* 04.2548 (13), and T6SS2 from *V. coralliilyticus* BAA-450 (14) were compared to the genomes of the 23 *Vibrio* isolates using CLICKER (https://cagecat.bioinformatics.nl/tools/clinker) (15). To identify MARTX toxins, we used the N-terminal region (1-2000 aa) from *V. vulnificus* strain 07-2444 RtxA protein (WP_158122697.1) to query the genomes of the *Vibrio* isolates using BLASTP (16). The results were filtered for >40% identity, E value <$10^{-5}$, and >70% coverage.

**GMT Island Analyses:** To identify GMT islands, the protein sequence of GmtY from *V. parahaemolyticus* RIMD 2210633 (WP_005477115.1) was used as a query in BLASTP against the genomes of the *Vibrio* isolates; results were filtered for >35% identity, E value <$10^{-5}$, and >70% coverage. To identify the 5′ and 3′ borders of each island, the nucleotide sequences 50 kb upstream and downstream of each *gmtY* gene were retrieved and used as a query in BLASTN against the RefSeq genome database. The resulting sequences from closely related *Vibrio* genomes were analyzed to identify those in which regions flanking the GMT island in the query genome are adjacent to each other. The islands' borders were then determined as previously described (17).

To identify T6SS toxins (effectors) within GMT islands, the sequences of the proteins encoded within the cargo of each island were used to identify known T6SS effector-associated domains, including PAAR, PAAR-like, Rhs, MIX, FIX, and RIX. As previously described (17). Predicted cognate immunity genes were identified as short genes found immediately downstream of predicted toxin genes.

Anti-phage defense systems encoded within the GMT islands were identified using the PADLOC (18) and DefenseFinder (19) tools. For PADLOC, GenBank DNA (.gb) sequences of the GMT islands were generated and provided as input. For DefenseFinder, amino acid sequences of GMT islands’ cargoes in FASTA format were used as input.

### Supplementary Figures

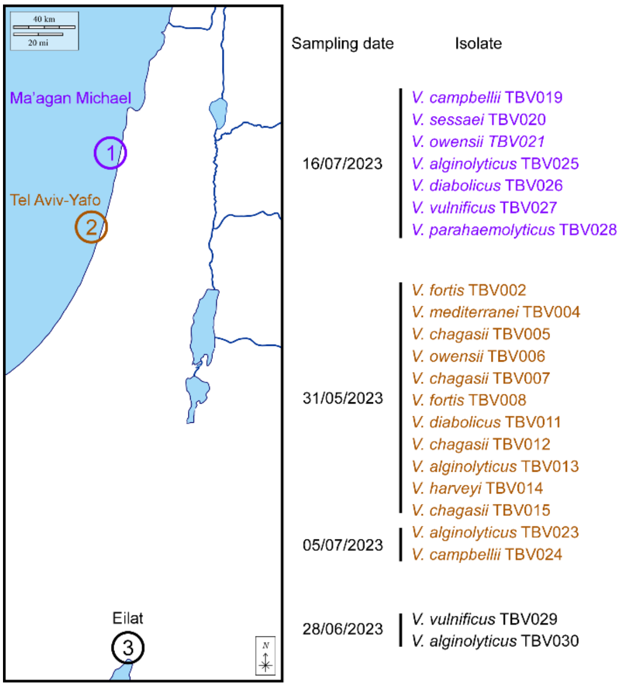


**Fig. S1. *Vibrio* sampling sites.** Sites 1 and 2 are in the Mediterranean Sea, and site 3 is in the Red Sea. Sampling dates and isolates recovered from each sample are denoted. The map was obtained from d-maps at https://d-maps.com/carte.php?num_car=338&lang=en.

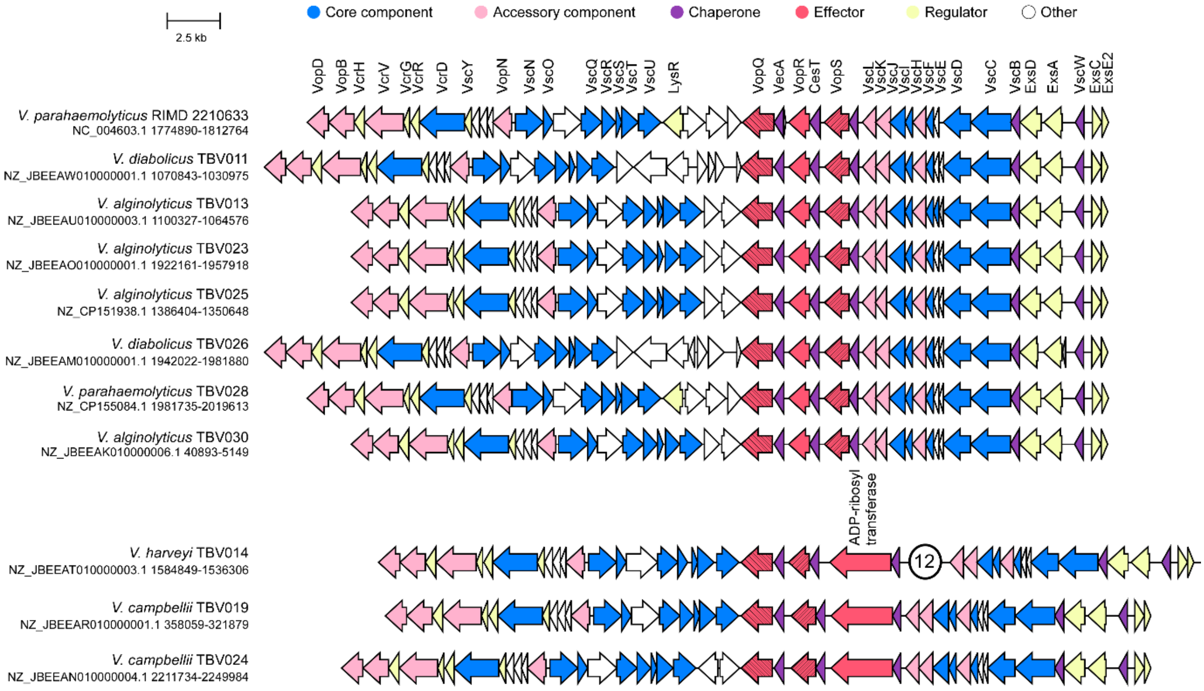


**Fig. S2. T3SS1-like gene clusters in environmental *Vibrio* isolates share a similar synteny.** Comparison of T3SS1 gene clusters found in the indicated *Vibrio* isolates and the representative T3SS1 gene cluster from *V. parahaemolyticus* RIMD 2210633. RefSeq accessions are denoted. Circled numbers denote the number of genes not shown.

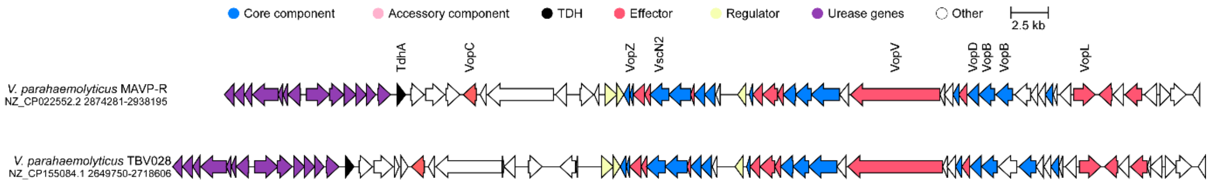


**Fig. S3. T3SS2-like gene clusters in environmental *Vibrio* isolates share a similar synteny.** Comparison of T3SS2 gene clusters found in the indicated *Vibrio* isolate and the representative T3SS2β gene cluster from *V. parahaemolyticus* MAVP-R. RefSeq accessions are denoted.

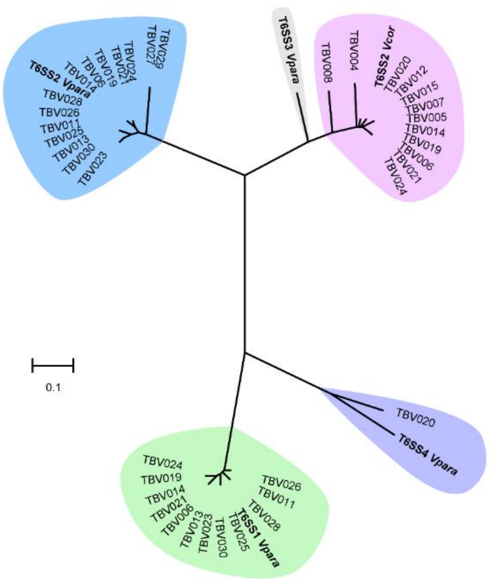

**Fig. S4. Four T6SS types identified in environmental *Vibrio* isolates.** Phylogenetic distribution of the T6SS core component TssB encoded within genomes of the environmental isolates, and representative TssB sequences for the different types of T6SSs known in vibrios (T6SS1-4 from *V. parahaemolyticus* [*Vpara*] and T6SS2 from *V. coralliilyticus* [*Vcor*]; shown in bold). The evolutionary history was inferred using the neighbor-joining method. The phylogenetic tree was drawn to scale, with branch lengths in the same units as the evolutionary distances used to infer the phylogenetic tree. The evolutionary distances were computed using the Poisson correction method; they are in the units of the number of amino acid substitutions per site.

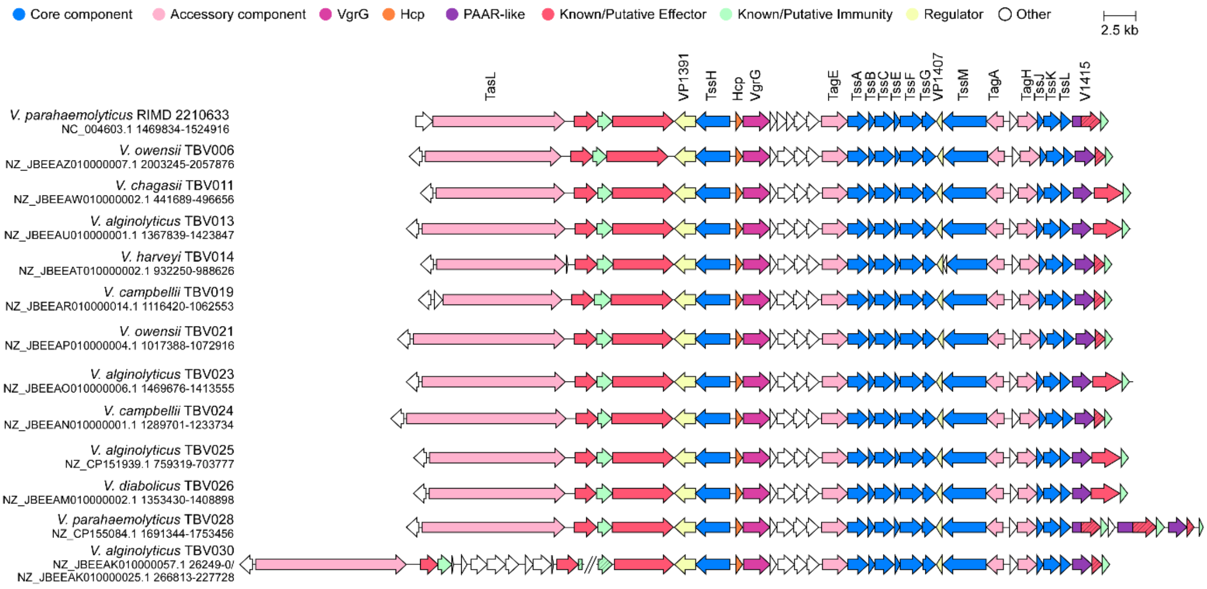


**Fig. S5. *V. parahaemolyticus* T6SS1-like gene clusters in environmental *Vibrio* isolates share a similar synteny.** Comparison of T6SS1 gene clusters found in the indicated *Vibrio* isolates and the representative T6SS1 gene cluster from *V. parahaemolyticus* RIMD 2210633. RefSeq accessions are denoted.

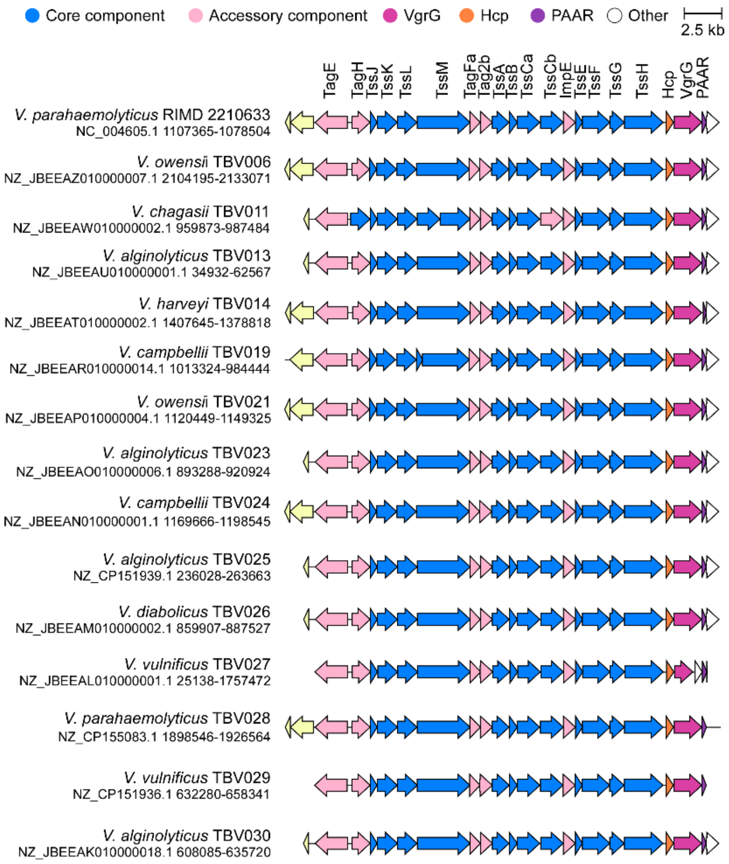


**Fig. S6. *V. parahaemolyticus* T6SS2-like gene clusters in environmental *Vibrio* isolates share a similar synteny.** Comparison of T6SS2 gene clusters found in the indicated *Vibrio* isolates and the representative T6SS2 gene cluster from *V. parahaemolyticus* RIMD 2210633. RefSeq accessions are denoted.

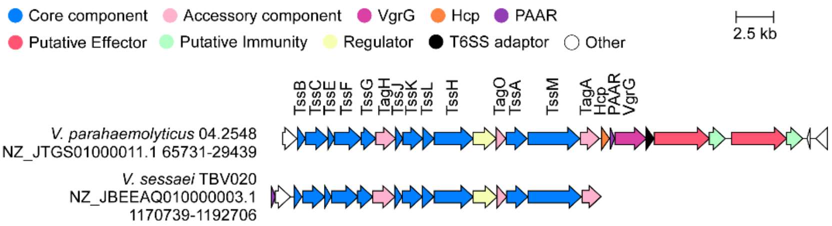


**Fig. S7. *V. parahaemolyticus* T6SS4-like gene clusters in environmental *Vibrio* isolates share a similar synteny.** Comparison of the T6SS4 gene clusters found in *V. sessaei* TBV020 and the representative T6SS2 gene cluster from *V. parahaemolyticus* 04.2548. RefSeq accessions are denoted.

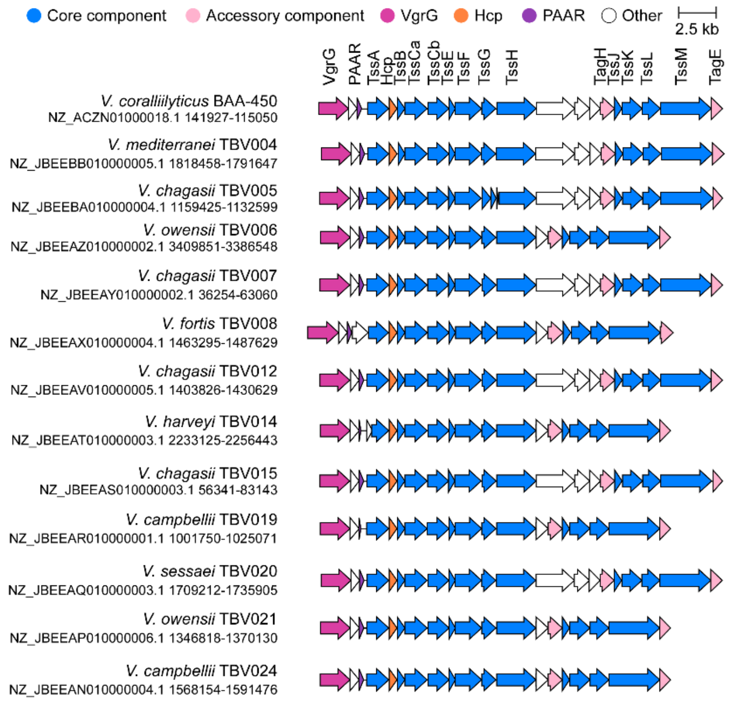


**Fig. S8. *V. coralliilyticus* T6SS2-like gene clusters in environmental *Vibrio* isolates share a similar synteny.** Comparison of T6SS2 gene clusters found in the indicated *Vibrio* isolates and the representative T6SS2 gene cluster from *V. coralliilyticus* BAA-450. RefSeq accessions are denoted.

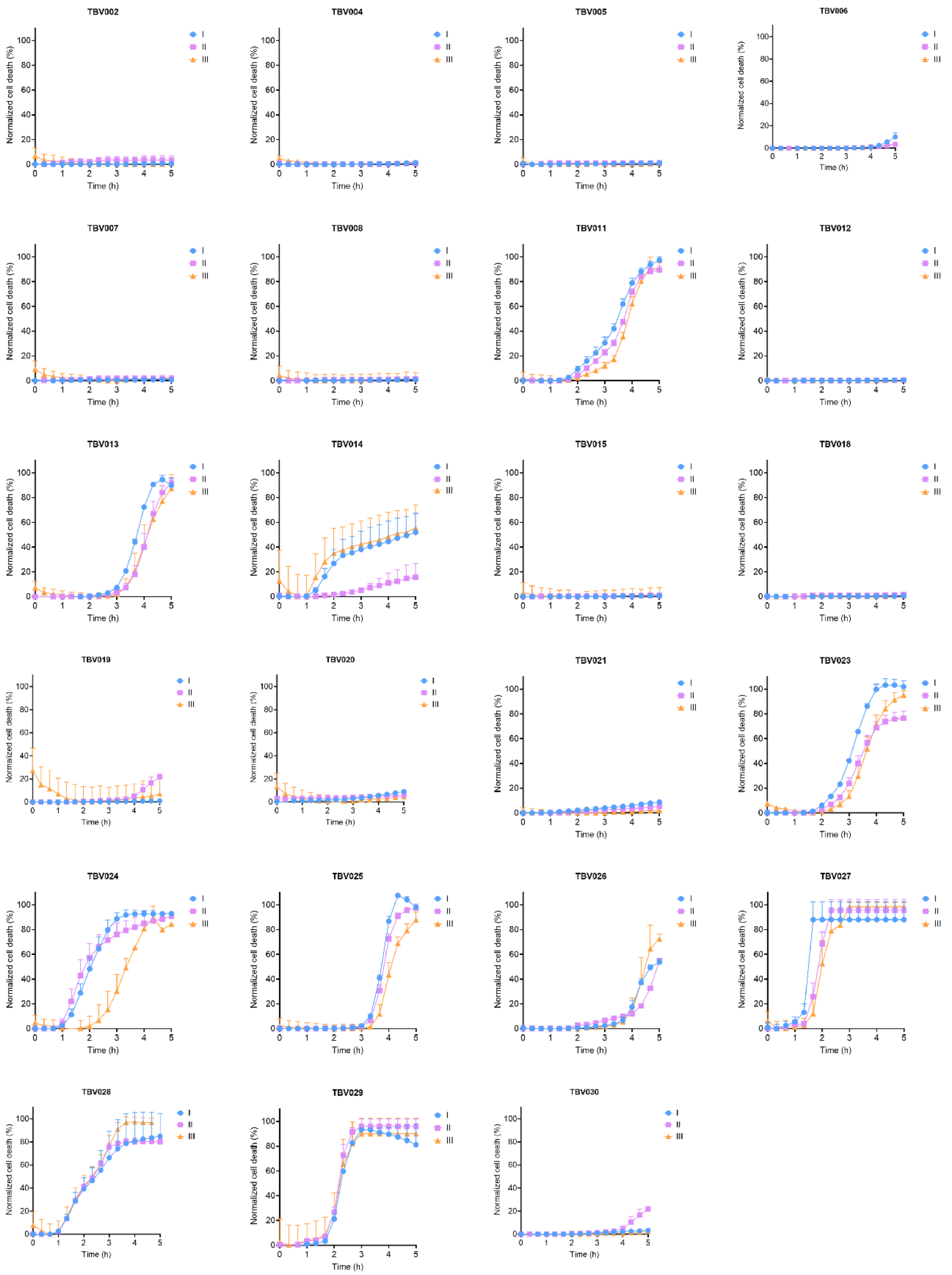


**Fig. S9. Environmental *Vibrio* isolates induce cell death in macrophages.** Cell death kinetics of bone marrow-derived macrophages (BMDMs) infected with the indicated environmental *Vibrio* isolates. The curves show 3 independent experiments (I, II, and III), each in a different color. Data are shown as mean ± SD, n = 3 independent samples. These curves were used to calculate the AUC summarized in Fig. 1B.

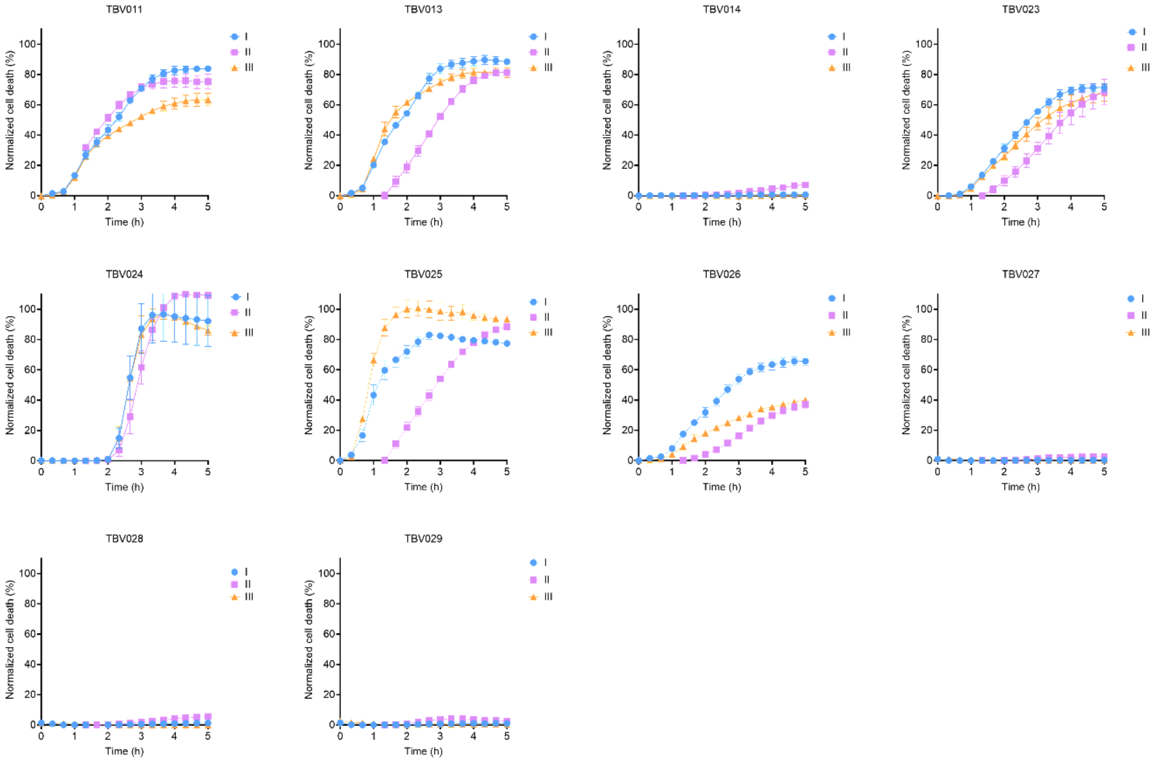


**Fig. S10. Spent media of environmental *Vibrio* isolates induce cell death in macrophages.** Cell death kinetics of bone marrow-derived macrophages (BMDMs) treated with cleared media from the indicated environmental *Vibrio* isolate cultures. The curves show 3 independent experiments (I, II, and III), each in a different color. Data are shown as mean ± SD, n = 3 independent samples. These curves were used to calculate the AUC summarized in Fig. 1C.

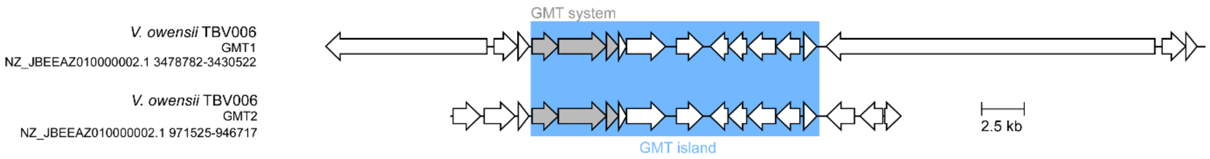


**Fig. S11. Identical GMT islands are found in two different locations on the genome of *V. owensii* TBV006.** The syntenies of two identical GMT islands found in *V. owensii* TBV006 (GMT_1 and GMT_2).

### Supplementary Tables

**Table S1. Information on *Vibrio* strains isolated in this study**

| Strain | Sampling location | Sampling date (DD/MM/YYYY) | GenBank assembly accession |
|---|---|---|---|
| *V. fortis* TBV002 | Tel Aviv, Israel | 31/05/2023 | GCA_046056525.1 |
| *V. mediterranei* TBV004 | Tel Aviv, Israel | 31/05/2023 | GCA_046057245.1 |
| *V. chagasii* TBV005 | Tel Aviv, Israel | 31/05/2023 | GCA_046057235.1 |
| *V. owensii* TBV006 | Tel Aviv, Israel | 31/05/2023 | GCA_046057115.1 |
| *V. chagasii* TBV007 | Tel Aviv, Israel | 31/05/2023 | GCA_046057095.1 |
| *V. fortis* TBV008 | Tel Aviv, Israel | 31/05/2023 | GCA_046057045.1 |
| *V. diabolicus* TBV011 | Tel Aviv, Israel | 31/05/2023 | GCA_046057055.1 |
| *V. chagasii* TBV012 | Tel Aviv, Israel | 31/05/2023 | GCA_046057035.1 |
| *V. alginolyticus* TBV013 | Tel Aviv, Israel | 31/05/2023 | GCA_046057005.1 |
| *V. harveyi* TBV014 | Tel Aviv, Israel | 31/05/2023 | GCA_046056995.1 |
| *V. chagasii* TBV015 | Tel Aviv, Israel | 31/05/2023 | GCA_046056975.1 |
| *V. chagasii* TBV018 | Tel Aviv, Israel | 31/05/2023 | GCA_046532705.1 |
| *V. campbellii* TBV019 | Ma’agan Michael, Israel | 16/07/2023 | GCA_046056945.1 |
| *V. sessaei* TBV020 | Ma’agan Michael, Israel | 16/07/2023 | GCA_046056895.1 |
| *V. owensii* TBV021 | Ma’agan Michael, Israel | 16/07/2023 | GCA_046056935.1 |
| *V. alginolyticus* TBV023 | Tel Aviv, Israel | 05/07/2023 | GCA_046056605.1 |
| *V. campbellii* TBV024 | Tel Aviv, Israel | 05/07/2023 | GCA_046056745.1 |

| | | | |
|---|---|---|---|
| *V. alginolyticus* TBV025 | Ma’agan Michael, Israel | 16/07/2023 | GCA_046532695.1 |
| *V. diabolicus* TBV026 | Ma’agan Michael, Israel | 16/07/2023 | GCA_046056635.1 |
| *V. vulnificus* TBV027 | Ma’agan Michael, Israel | 16/07/2023 | GCA_046056645.1 |
| *V. parahaemolyticus* TBV028 | Ma’agan Michael, Israel | 16/07/2023 | GCA_046532685.1 |
| *V. vulnificus* TBV029 | Eilat, Israel | 28/06/2023 | GCA_046532655.1 |
| *V. alginolyticus* TBV030 | Eilat, Israel | 28/06/2023 | GCA_046056815.1 |

### Supplementary Datasets (caption)

**Dataset S1. TssB sequences used to construct a phylogenetic tree.**

**Dataset S2. Antibiotics MIC values from three independent experiments.**
